# MSCProfiler: An image processing workflow to investigate Mesenchymal Stem Cell heterogeneity using imaging flow cytometry data

**DOI:** 10.1101/2023.05.13.540536

**Authors:** A. Gupta, S.K. Shaik, L. Balasubramanian, U Chakraborty

## Abstract

Single-cell immuno-heterogeneity has always been the forerunner of any change in homeostasis of cellular functions in the body. Mesenchymal stem cells represent a viable source for the development of cell-based therapies. Multiple conditions giving rise to inter, and intra-population variations result in heterogeneity and multipotent differentiation ability of these cells of stromal origin. Cell surface markers which are important members of membrane proteins, ion channels, transporter, adhesion, and signaling molecules generally differentiate between stromal cells of different origin. However, existing analytical tools cannot always model a pattern of their surface distribution in successive generations of growth and proliferation. In this study, we have developed a post-acquisition image analysis pipeline for human mesenchymal stromal cells obtained from exfoliated deciduous teeth (hSHEDs). Using the open-source image processing software CellProfiler, a pipeline has been developed to extract cellular features from 50,000-100,000 single-cell images. We made use of the image flow cytometry technology to explore the morphometric properties of hSHEDs, along with their surface marker distribution. This unbiased pipeline can extract cellular, geometrical and texture features such as shape, size, eccentricity, entropy, intensities as a measure of cellular heterogeneity. For the first time, we have described an automated, unbiased image assessment protocol implemented in a validated open-source software, leveraging the suite of image-based measurements to develop the prototype named as MSCProfiler. The hallmark of this screening workflow has been the identification and removal of image-based aberrations to identify the single-cell bright field and fluorescent images of mesenchymal stem cells.

## INTRODUCTION

At the forefront of cell-based therapies are Mesenchymal Stem Cells (MSCs) with over 1000 registered clinical studies underway (1). These are adult tissue derived multipotent stem cells which can be isolated from almost all post-natal organs and tissues (2). They are known to display a variety of pleiotropic functions such as anti-apoptosis, angiogenesis, anti-fibrosis, and chemo-attractive properties, making them a useful tool for a wide variety of therapeutic applications (1,3,4). An important metric to qualify MSCs as beneficial therapeutic agent is the determination of their safety and risks, if any (4). Consequently, culturing and studying these cells become important. However, one of the hurdles faced in the translation of *in-vitro* MSC studies is that many of the properties that are studied arise because of the artificial culture environment (5). Expression of surface markers such as CD90, CD73 and CD105, which are routinely used for MSC characterization, are limited to only being expressed *in-vitro* and not *in-situ*. Additionally, differentiation ability of MSCs into mesoderm lineages has also not been proven to happen *in-vivo* (5,6). A major challenge in the translation of MSCs to clinical application is the inherent heterogeneity of MSCs which influences its properties of immunomodulation and regeneration(3). The heterogeneity, arising due to isolation, culture, and expansion, warrants an extensive study on various MSC subtypes and their categorization (3,7).

For us to understand this heterogeneity, a cell-by-cell approach as opposed to conducting studies at a population level would be more advantageous in revealing important and unique properties of various cell types (8). Single-cell analysis platforms such as imaging flow cytometry (IFC) can prove to be a powerful tool. An amalgamation of high-throughput cytometry and microscopy, IFC allows extraction of traditional flow cytometric data and images of each event from fluorescent, brightfield (BF), and laser side scatter/ darkfield (DF) channels (9). This technology has allowed scientists to perform advanced assays such as nuclear translocation (10), autophagy (11,12), detection of DNA damage (13,14), and cell division (15), to name a few. The high-throughput nature of this technology allows collection of enormous amounts of data from a single cell which can be used to answer questions of cellular heterogeneity (9) and holds immense potential for data mining application and creating well trained neural networks. However, the large amount of information remains largely under-utilized due to the lack of data analysis tools that can extract meaningful information from the images (9).

The Amnis Imagestream image cytometers are one of the commonly used IFC instruments. Data analysis on Amnis platforms can be done using their proprietary software IDEAS or with analysis tools such as FCS Express (DeNovo Software). Apart from having the advantage of performing traditional flow cytometry data analysis, the IDEAS platform also provides users with ‘wizards’, which provide assay specific analysis templates such as apoptosis, colocalization, shape change, feature finder, spot count, that enable sequential analysis in general without many complications (https://cytekbio.com/pages/imagestream). Additionally, specific areas or regions of the cell image can be defined using the ‘masking’ (defined set of pixels in the region of interest) that contain feature such as creating nuclear masks or cell surface marker specific masks. It is also possible to create the masks on IDEAS based on user-defined criteria. This can be done by using the mask manager which has 13 available functions to create the new mask. However, use of this analytical software is primarily dependent on the experience of the user and therefore brings in bias. This can especially be challenging when understanding immuno-heterogeneity which can lead to missing out on potentially important features if one does not actively look for them.

The recent trend in the incorporation of machine learning (ML) or artificial intelligence (AI) in biology has resulted in advancement of data analysis approaches (16). Use of virtual or label-free staining of cells (17,18), use of brightfield (BF) information alone to extract quantitative features of cells (17), and use of ML to analyse IFC data for diagnostics (19) have all introduced a new paradigm. However, some of these methods might require an in-depth knowledge of deep-learning techniques, neural networks, or programming languages. In this study, we have used a user friendly, freely available software CellProfiler (available at https://cellprofiler.org/) to create a pipeline that can be used to analyse most of the commonly available image file formats (20).

Here we introduce MSCProfiler, a CellProfiler pipeline for the analysis of IFC data of MSCs. Stem cells derived from human exfoliated deciduous teeth (hSHEDs) have been stained with antibodies which recognize surface antigens of these cells, prescribed as minimal criteria for identification of MSCs by the International Society for Cell & Gene Therapy (ISCT) (6). We have included two surface markers in our studies-CD44/ homing cell adhesion molecule (HCAM) and CD73/ ecto-5’-nucleotidase. CD44 is a glycoprotein with its primary function being related to cell attachment and migration. It is involved in lymphocyte homing, hyaluronate degeneration, and functions as a transcription mediator. In MSCs, CD44 interacts with hyaluronan present in the surrounding environment to facilitate migration (21). CD73 is a glycosylphosphatidylinositol (GPI) anchored glycoprotein and has many roles such as cell adhesion, generation of adenosine, and has also been implicated in mediating anti-inflammatory role in MSCs (22). Conventionally, with routine flow cytometry based characterization of MSCs, bivariate plots of CD44 and CD73 should represent ≤95% double positive population to suffice the criteria of MSCs as per ISCT guidelines and should express ≥2 % hematopoietic marker CD45 (6). In our panel we have used the above markers and also included Sytox Green as a live/dead discriminator. Live cells exclude Sytox Green from their membranes, whereas the nucleus of dying/dead cells whose membrane integrity have been compromised take up the dye (23). We have focused on extraction of feature information from individual BF and/or fluorescent channels/ detectors available on the Amnis ImageStream Mk II instrument. This pipeline can identify live singlets based on the exact boundary of the cells from the BF images of MSCs; compute the geometrical parameters of the singlets such as area, aspect ratio, eccentricity, compactness, and even texture features like inverse difference moment, and entropy. The intensity of CD44 and CD73 antigens were also extracted from the fluorescent channel images and their surface distribution pattern was observed.

## MATERIALS AND METHODS

### Cell Culture

hSHEDs cultured at passage 9 were seeded in 100mm culture dishes at a density of 5000 per cm^2^ and grown in KnockOut DMEM (Gibco) media supplemented with 10% FBS (Gibco), 1X Pen-Strep (Gibco) and 1X Glutamine (Gibco) and grown at 37°C (humidity conditions) until they reached 90-95% confluency with media changes every 24 or 48 hours as required. Cells were then washed once with plain basal media without any supplements and trypsinization was done by adding 0.5% Trypsin-EDTA (Gibco) followed by incubation at 37°C for 10 minutes until all the cells had detached from the culture dish. Trypsin-EDTA was neutralized with basal media and the cells were collected and centrifuged at 1200 rpm for 6 minutes to obtain the cell pellet.

### Preparation of cell suspension for immunostaining

Cell pellet was resuspended in 1mL media, and the cell count was determined. Cells were then washed twice with staining buffer (2% FBS in PBS). 2% FBS helps maintain live cells while in suspension. A cell suspension of 1 × 10^6^ cells per 50 µL suspension, contributed to 100µL of reaction volume when mixed with antibodies and staining buffer. The amount of antibody added to every tube has been described in Table 1.

**Table 1:**
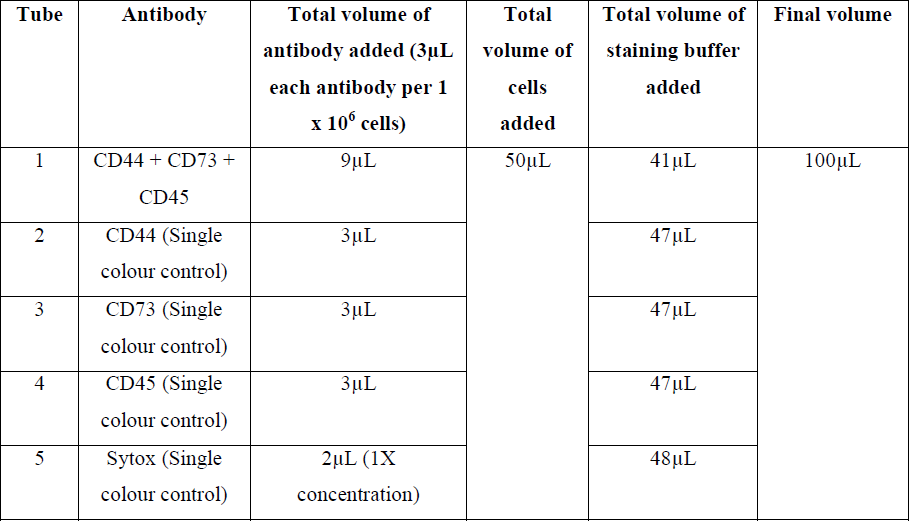
Sample Preparation Guide

| **Tube** | **Antibody** | **Total volume of antibody added (3μL each antibody per $1 \times 10^6$ cells)** | **Total volume of cells added** | **Total volume of staining buffer added** | **Final volume** |
| --- | --- | --- | --- | --- | --- |
| 1 | CD44 + CD73 + CD45 | 9μL | 50μL | 41μL | 100μL |
| 2 | CD44 (Single colour control) | 3μL | | 47μL | |
| 3 | CD73 (Single colour control) | 3μL | | 47μL | |
| 4 | CD45 (Single colour control) | 3μL | | 47μL | |
| 5 | Sytox (Single colour control) | 2μL (1X concentration) | | 48μL | |

### Antibodies and immunostaining

Live hSHEDs were simultaneously stained for 3 surface antigens with an antibody cocktail consisting of CD44, CD73 and CD45. The antibody-fluorochrome conjugates have been described in Table 2. Cells were incubated for 30 minutes and washed twice with staining buffer and finally resuspended in 100uL of staining buffer. Prior to acquisition, cells were stained for 15 minutes at 37°C with SYTOX Green (1X) was added to tube 1, a nucleic acid stain to distinguish dead cells from live cells.

**Table 2:** Antibody-fluorochrome conjugates

| ANTIBODY | FLUOROCHROME TAG | COMPANY | EXCITATION MAXIMA | EMISSION MAXIMA | EXCITATION LASER |
| --- | --- | --- | --- | --- | --- |
| Anti-CD73<br>(Human) | PerCP-Cy5.5 Mouse<br>Anti-Human | BD<br>Biosciences | 482nm | 690nm | 488nm |
| Anti-CD44<br>(Human) | BD Horizon V450<br>Mouse Anti-Human | BD<br>Biosciences | 404nm | 448nm | 405nm |
| Anti-CD45<br>(Human) | PE Mouse Anti-Human | BD<br>Biosciences | 566nm | 574nm | 488nm |

To address the issue of spectral spillage by applying the compensation algorithm, single colour sample tubes for each antibody were also prepared. For this, only a single antibody per tube, along with cell suspension (MSCs) was added. Since MSCs do not express CD45, to obtain the single colour data, whole blood was collected and stained for CD45 (pan leukocyte marker). To obtain single colour control files for SYTOX Green, cells were given heat shock treatment to ensure majority of the cells would be non-viable and thus take up the dye. Cells were kept in 70°C for 5 minutes and the immediately placed on ice. The reaction mix in the subsequent tubes has been summarized in Table 1.

### Acquisition on the Amnis ImageStream Mk II

Cells were acquired on the IFC platform Amnis ImageStream Mk II equipped with 1 charged couple device (CCD) camera (6 channels/ detectors) using the INSPIRE acquisition software, briefly described in Fig. 1(a). The detection channels available for use in the instrument based on our fluorochrome conjugates were - V450 (Channel 1, 435-505nm), SYTOX Green (Channel 2, 505-560nm), PE (Channel 3, 560-595nm) and PerCP-Cy5.5 (Channel 5, 642-745nm), brightfield (BF) images (Channel 4, 595-642nm) and darkfield images (DF)/ Side scatter (SSC) (Channel 6, 745-780nm). Table 3 includes the instrument configuration and panel design. Unstained cells were first run to set the baseline correction by adjusting the laser powers such that no autofluorescence could be detected. Lasers were set at 10mW for 405nm laser (Channel 1 excitation) and 10mW for 488nm laser (Channel 2, 3 and 5 excitation). Sample tubes containing all the 4 colours were run and 50,000 to 1,00,000 events were acquired. All the single-colour tubes were run with the BF and DF parameters turned off, and automatically got saved with ‘noBF’ mentioned in their file names. These files were later used to set up the compensation matrix post-acquisition. All raw data files were saved in raw image file (.rif) format.

**Fig. 1:**
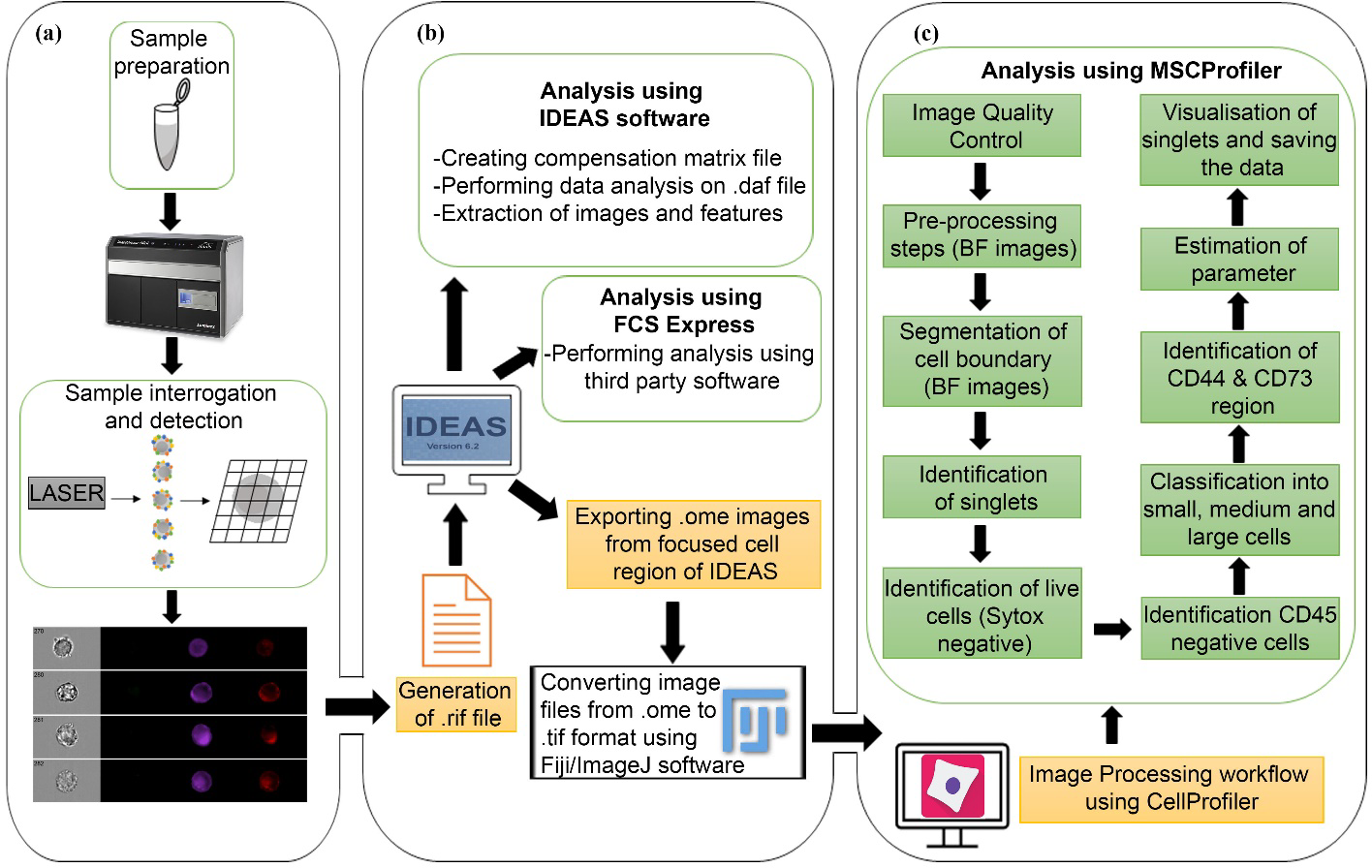
Supervised versus automated data analytical tools. **(a)** Sample preparation and acquisition on the imaging flow cytometer using the INSPIRE software, which generated the single cell images of each cell in flow. **(b)** Supervised analytical software IDEAS v6.2 was used to generate compensation matrix and .daf and .cif files to analyse the acquired data and view the cell images. Data analysis software such as FCS Express v6 was used to analyse the cytometry data. **(c)** MSCProfiler analytical automated pipeline. Images exported from IDEAS were converted to .tiff format and later uploaded on MSCProfiler and run through the pipeline to extract information from the single-cell images and identify cells of interest.

**Table 3:**
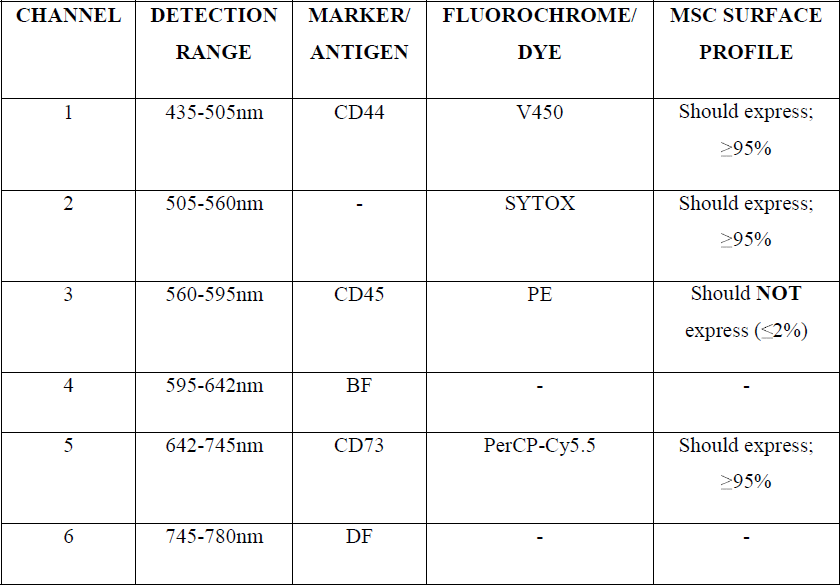
Amnis ImageStream Mk II configuration and panel design.

### Setting up compensation on IDEAS

As opposed to conventional flow cytometry that relies on voltage gain of individual photomultiplier tubes (PMTs) to collect signal generated from antibody fluorochrome conjugate, the detectors available on the Amnis ImageStream Mk II are CCDs. They have a higher quantum efficiency than PMTs which make them ideal detectors to collect dim fluorescence signals a prerequisite for good imaging. On this instrument, a pixel-based compensation is performed post-acquisition using the IDEAS software. The ‘noBF’ files of each colour acquired in .rif format were loaded onto the IDEAS software to create a compensation matrix to correct the spectral spillage of one fluorochrome to other channels. Once the compensation matrix is set up, they get saved in compensation matrix (.ctm) file format.

### IFC data analysis

The IFC data has two components-first, the flow cytometric data conventionally represented as dots on a plot and second, the individual images of each dot on the plot. The data analysis was performed using the proprietary software IDEAS v6.2 (Amnis Corp, Seattle, WA). Following a Boolean logic, the gated population of interest were first identified after which the image data analysis begins. Every cell is demarcated with a number and is visible on the screen/ image gallery. ‘noBF’ files were used to set up the compensation matrix. The .rif and .ctm files were then used to create the data analysis file (.daf) and compensated image file (.cif). The downstream analysis was performed using the .daf files. Following a hierarchical gating strategy, first the focused cells were identified by plotting a histogram of normalized frequency versus gradient root mean square (RMS) value of Channel 4 (BF). This was followed by single cell identification (selected apart from the aggregates/ debris) gated from the aspect ratio of Channel 4 versus area of Channel 4. The next plot was to identify live cells (negative for Sytox) from the Channel 6 (DF) vs Channel 2 plot, followed by CD45 negative cells from the Channel 6 (DF) vs Channel 3 (CD45) plot (exclusion gating strategy) and finally populations positive for both CD44 and CD73 were identified from a bivariate plot between Channel 1 (CD44) and Channel 5 (CD73).

### FCS Express Analysis

The .daf file was read by the FCS Express v6 (Research Use Only), a licensed flow cytometry data analysis software. It was ensured that both the .daf file and the .cif file for each sample tube was available in the same analysis folder. Using similar gating strategy described in the preceding section, singlets were identified first, followed by live cells. The CD45 negative cells were first identified and gated onto the next plot to show the CD44 and CD73 double positive cells. On this platform, image data could be viewed only and not analyzed. A flow chart of the steps involved in supervised analysis platforms have been shown in Fig. 1(b).

### Preparing images for CellProfiler Analysis

The individual images from all channels 1, 2, 3, 4 and 5 were selected from the ‘Export .tif images’ under the ‘Tools’ option on IDEAS. The images come with open microscopy environment (.ome) extension which were converted to the .tiff using the Fiji/ImageJ software in an automated fashion using a macro (supplementary information). These individual images were used in the image processing workflow as described in the later sections, to extract the single cellular features. The workflow was developed using the CellProfiler software to analyse the single-cell images of MSCs.

### MSCProfiler Pipeline

The individual .tiff images of all the channels were then loaded into MSCProfiler pipeline through the input modules (features that perform specific tasks). Using the ‘NamesAndTypes’ module each channel image has been categorized in a way that a singlet set consists of information on a single cell/ event consisting of BF and fluorescent channel images. The automated pipeline has been described by broadly classifying them into 11 steps based on the purpose they serve. Individual cell information, i.e., information from all 5 channels, is processed through all the modules of each component to identify and extract various features like geometric, texture and intensity values, described in Fig. 1(c). The total number of cells analyzed was 54,356, which were selected from the ‘focused’ cell gate on the IDEAS software. Therefore, a total of 271,780 images, collected from all five channels, were run on MSCProfiler. Due to the limited computational power all the 271,780 images were not run through the workflow at once time. The total set has been divided into 24 groups each containing approximately 10,000 images. The steps in the MSCProfiler workflow are as follows:

1. Quality Control of Images – The MeasureImageIntensity and FlagImage modules were used to discard images that do not contain any cellular information. The MeasureImageIntensity module calculated the intensity measurements of the images based on which the boundary conditions that were set in the FlagImage module to discard the poor-quality images. Only BF (Channel 4) images were used for this quality check to estimate if the image has any cellular information or not. Those BF images whose total image intensity was in the range of 5,500-7,900 pixels and whose total image area was in the range of 9,000-13,000 pixels were best fit to proceed. The images that did not satisfy the criteria were discarded/flagged from the analysis.
2. Pre-Processing Steps – The unflagged image set was then used in further analysis. To identify the cellular region BF images were used. Each of the BF images were pre-processed using four modules ImageMath, Smooth, RescaleIntensity and EnhanceEdges. The ImageMath module was used for subtraction of the foreground from the background of the BF images. The Smooth module applied the Gaussian filter on the BF images to blur the pixels outside the cell and highlight the pixels inside the cell. The RescaleIntensity module found the minimum and maximum intensity values across the entire BF image and rescaled every pixel so that the minimum intensity value was zero and the maximum intensity value was one. The EnhanceEdges module made use of the Sobel filter method to highlight the edges of the cell identified in the BF image.
3. Segmentation of Cell Boundary - The pre-processed BF images resulting from the above-mentioned steps were used to identify the cell boundary. This section of the pipeline involved four modules, first, the threshold module was used to identify the pixels of interest based on the grey scale signal using the Otsu thresholding, a method based on the signal variances between the foreground and the background. Second, the Morph module removed the identified pixels from the background of the segmented image. Third, IdentifyPrimaryObjects module identified cell boundary. Lastly, ExpandOrShrinkObjects module was added to get a more accurate cellular boundary. The BF images which contained one cell per image proceed for further processing while those which contained more than one cell per image were discarded by the FlagImage module.
4. Identification of Singlets – This section of the pipeline contained three modules that were used to identify only the singlets MSCs and remove the images that contained more than one cell. The MeasureObjectSizeShape module calculated geometrical features like Area, FormFactor, Compactness, Eccentricity, of the cell identified from the BF image. A boundary condition was set up on the Area, FormFactor and Eccentricity measurements in the FilterObjects module to obtain the singlets. Eccentricity describes how deviated a shape/ object is from a circular shape, for a circle it is equal to 0. Compactness gives us an idea if the shape/ object has a hole (or missing pixels) in it or is close bound figure, a circle has a compactness of 1. Aspect ratio (minimum ferret diameter/ maximum ferret diameter) defines how elongated an object is, for a perfect circular object it is equal to 1. For the identified cell to be a singlet its Area was set in the range of 150-4500 pixels, its FormFactor lied in the range of 0.75-1.0 pixels and its Eccentricity fell in the range of 0.0-0.7 pixels. If the identified cell did not meet this criterion, the BF image was considered as a non-singlet and is discarded by the FlagImage module. These ranges described throughout the were set by first taking a set of 50-100 cell images that had mix of target and non-target cells and were measured during the pipeline building steps.
5. Identification of Live Singlets – This part of the pipeline confirmed an identified singlet as a live or a dead cell based on the appropriate marker using the following five modules-RescaleIntensity, MeasureObjectIntensity, FilterObjects, MeasureImageIntensity and FlagImage modules. These modules sequentially called for the Sytox channel images (Channel 2) of the identified singlet. For a singlet to be a live singlet, the mean intensity value of the Sytox channel was set to 0 pixels. Therefore, such cells with no signal in the Sytox channel were the live cells. Detection of signal/ intensity from the Sytox channel, marked the cell as a dead cell and hence discarded.
6. Identification of CD45 Negative Population – This section of the pipeline checked that the MSCs are negative for CD45 (tagged to PE) as one of the first criteria to identify them. The RescaleIntensity, MeasureObjectIntensity, FilterObjects, MeasureImageIntensity and FlagImage modules were again sequentially called for the PE channel (Channel 3) images of the identified live singlets to confirm that the cells were PE negative. The mean intensity value of the PE channel image lied in the range 0-0.002 pixels.
7. Classification of Singlets into Large, Medium, and Small MSCs – The identified singlets of MSCs were categorized as small, medium and large cells based on their area and this was performed using the FilterObjects module. The small, medium, and large MSC singlet classification was based on the area criteria set within the range of 2200-3000, 3001-4000 and 4001-4500 pixels respectively.
8. Identifying CD44 and CD73 Surface Marker Regions – The region of expression of CD44 (Channel 1) and CD73 (Channel 5) were identified using the Identify Primary Object module based on the minimum cross entropy thresholding method.
9. Estimation of Parameters: There are various parametric features that were calculated for the identified singlets based on the geometry, texture, and intensity measurements. The modules include Measure Texture and Measure Granularity. Using the BF images, a bunch of features like area, eccentricity, compactness, entropy, granularity, were calculated. Using the identified CD44 and CD73 regions the total intensity of both the markers were also measured. Texture Features like inverse difference moment and entropy value were also calculated. Inverse difference moment gave a measure of the homogeneity within a defined region. Entropy value indicated the randomness within the structure.
10. Visualization of Singlets – The OverlayOutline module gave the outline of the cell and outline of the CD44 and CD73 markers on the BF image of the cell. These images were saved as .png files with the help of the SaveImages module.
11. Saving data – In the last section of the pipeline, the data of all the calculated values were saved as separate .csv files with the help of the ExportToSpreadsheet module. The data of the geometrical features of the singlets were also saved as a .properties file by the ExportToDatabase module. This .properties file can be used in future for analysis and classification of the MSCs using the ML approach. An overview of the methods has been illustrated in Fig. 1 from the acquisition to the analysis stage.

## RESULTS

### Conventional flow cytometry-based analysis versus imaging cytometry analysis

The workflow followed in this study, shown in Fig. 1 schematic, compares supervised analytical tools with our automated data analysis pipeline, MSCProfiler. The IFC results were analyzed on the FCS Express software following the gating strategy as shown in Fig. 2(a). The next set of analysis following similar flow cytometric logic as shown in Fig. 2(b), was performed using IDEAS. The FCS Express analysis revealed that out of 142,171 (the total acquired events) the number of singlets were 10,314, out of which 9,733 were live cells, i.e., negative for Sytox Green, 9,714 were CD45 negative and 9,404 cells were double positive for both CD44 and CD73, shown in Fig. 2(a). Similarly, using the IDEAS software, out of 142,171 total events, 54,356 were focused cells. From the focused cells, 7,900 cells were identified as singlets, 7,700 were live cells. Lastly, the number of cells positive for both CD44 and CD73 were found to be 7,579, as seen in Fig. 2(b). The gate statistics and the percentages for both the software have been summarized in a tabular format in Fig. 2(c). The sequential output of IDEAS gating strategy has been described in Fig. 3(a). The visual confirmation of every acquired event during the analysis stage, through the image gallery of IDEAS, serves to bolster the gating strategy on the platform. For example, from the image gallery the lives cells were confirmed by comparing against the images of the cells that had taken up Sytox Green and showed nuclear localization, versus the absence of any signal in the channel 2 for the live cells as shown in Fig. 3(a).

**Fig 2:**
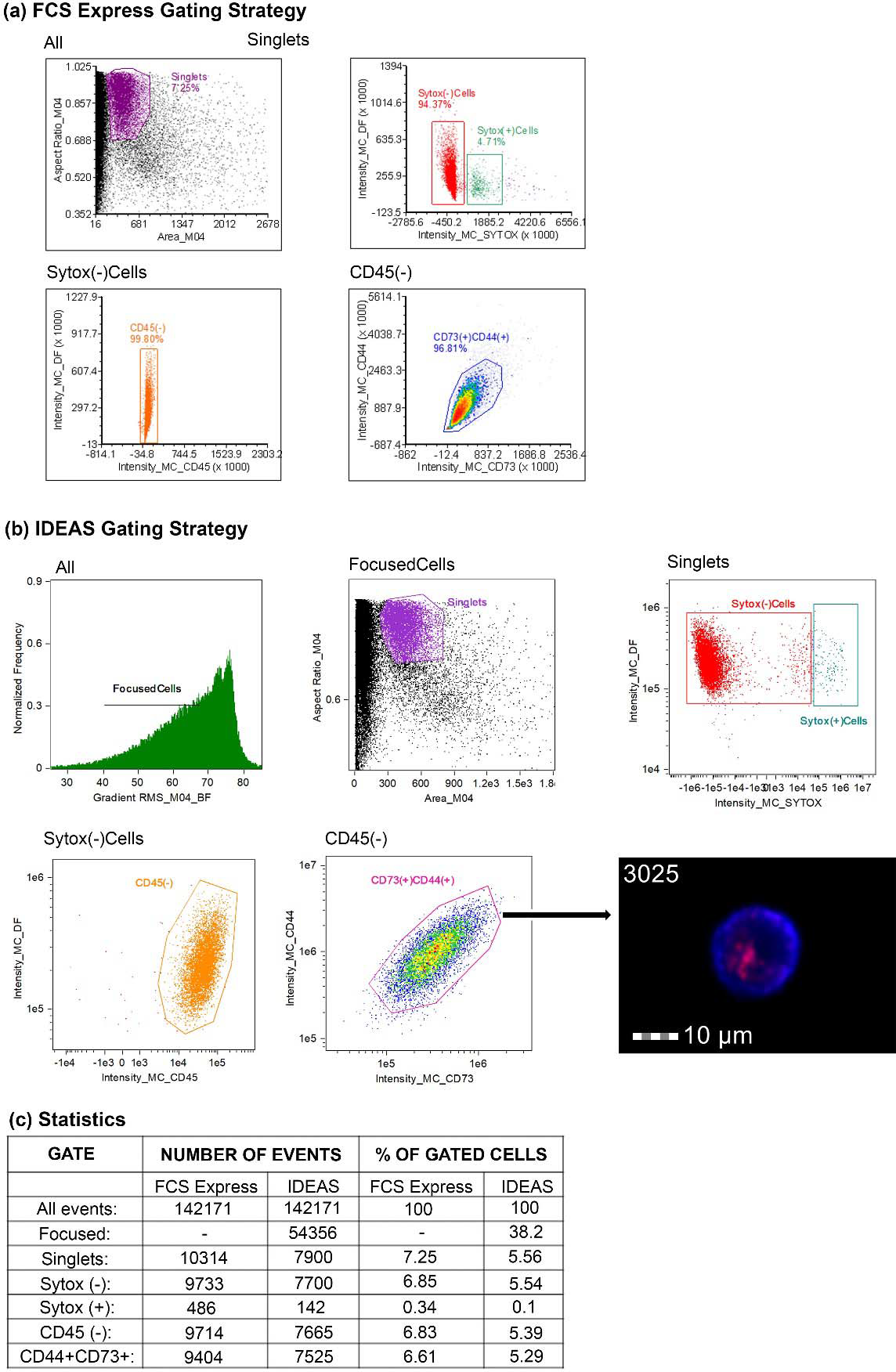
A comparison between the software FCS Express and IDEAS with regard to their gating strategies used. **(a)** FCS Express gating strategy show a hierarchical approach to identify target cells which first starts with gating singlets, followed by identification of live cells (negative for Sytox Green, followed by exclusion gating of CD45 negative cells and then identification of double positive populations of CD73 and CD44 expressing cells. Gate hierarchy and statistics have been indicated in the figure. **(b)** IDEAS gating strategy followed a similar sequential gating strategy that first identified focused cells, followed by the identification of singlets. Here also the same gating hierarchy was followed as above to identify the double positive expressors (CD44^+^ CD73^+^ MSCs). A representative image of a single live hSHED (Cell no. 3025) from the image gallery on the Amnis Imagestream shows the double expressor of CD44 and CD73 **(c)** The statistics of the different populations analyzed by these supervised platforms have been tabulated.

**Fig 3:**
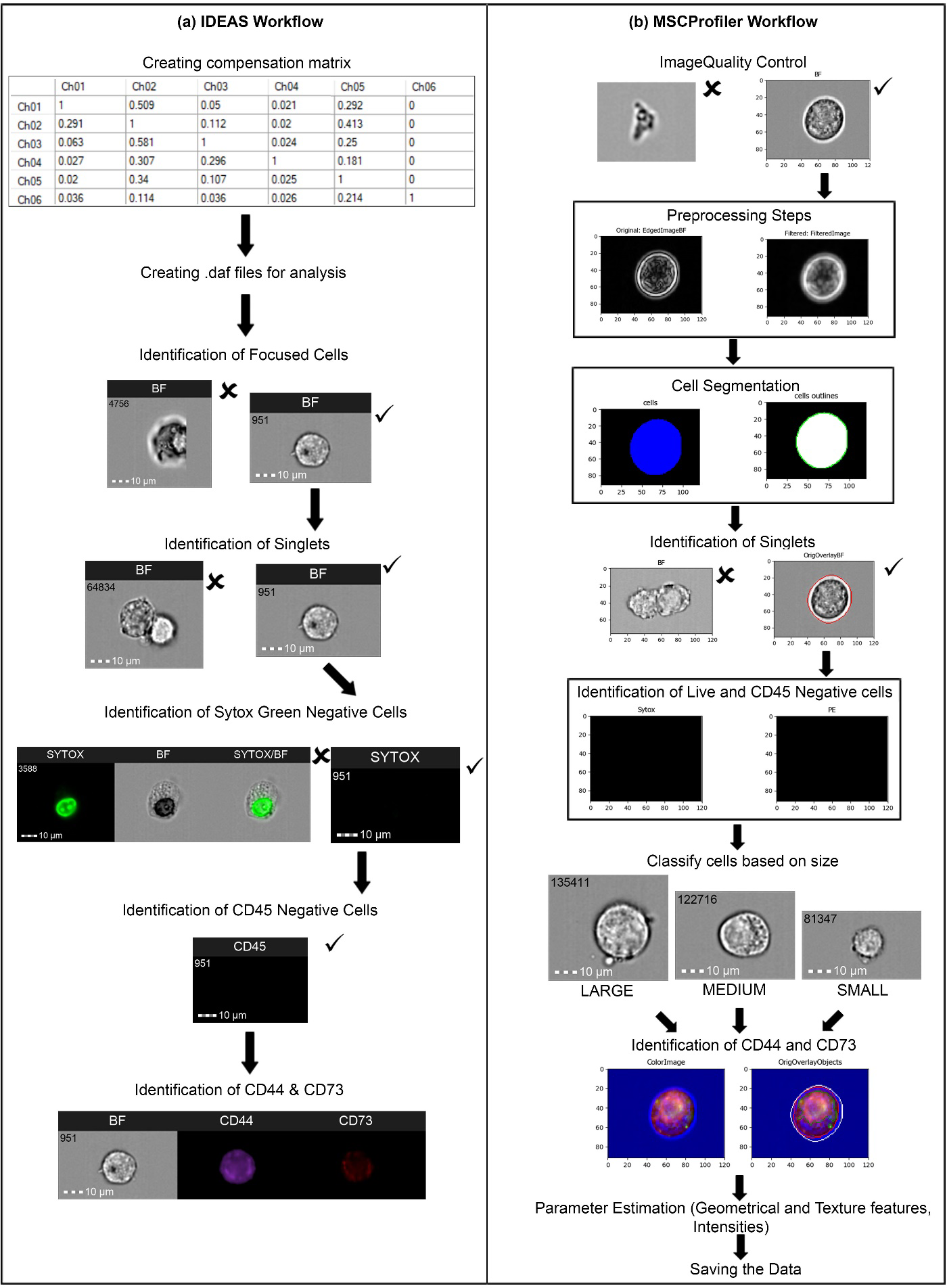
Stepwise identification of populations of hSHEDs using the supervised and our automated workflow, MSCProfiler. **(a)** Sequential output of IDEAS workflow following the gating strategy as described. Representative cell images in each step have been shown here. **(b)** Sequential output of MSCProfiler workflow shows a comparable end result. However, in the first case, data input includes the .rif, while in MSCProfiler the data input is in the form of single cell images.

In Fig. 4, single cell images as seen on the IDEAS image gallery has been shown. In Fig. 4(a), the width of each cell was also calculated from the BF channel image (top right corner of each image). We observed that the cells varied in their sizes and manually segregated them into small cells had a width of 16 to 20, medium cells were 20 to 26 and large cells were 27 and above. In Fig. 4(b), the CD44 and CD73 double positive expressors have been shown. A lack of signal can be seen from the Sytox and CD45 channels. Additionally, to understand the dual expression pattern of the two surface markers we used the IDEAS masking feature, CD44 and CD73 staining which have been represented by pseudo colours blue and red, respectively, showed a pink to purple tinge in areas where dual expression could be seen.

**Fig 4:**
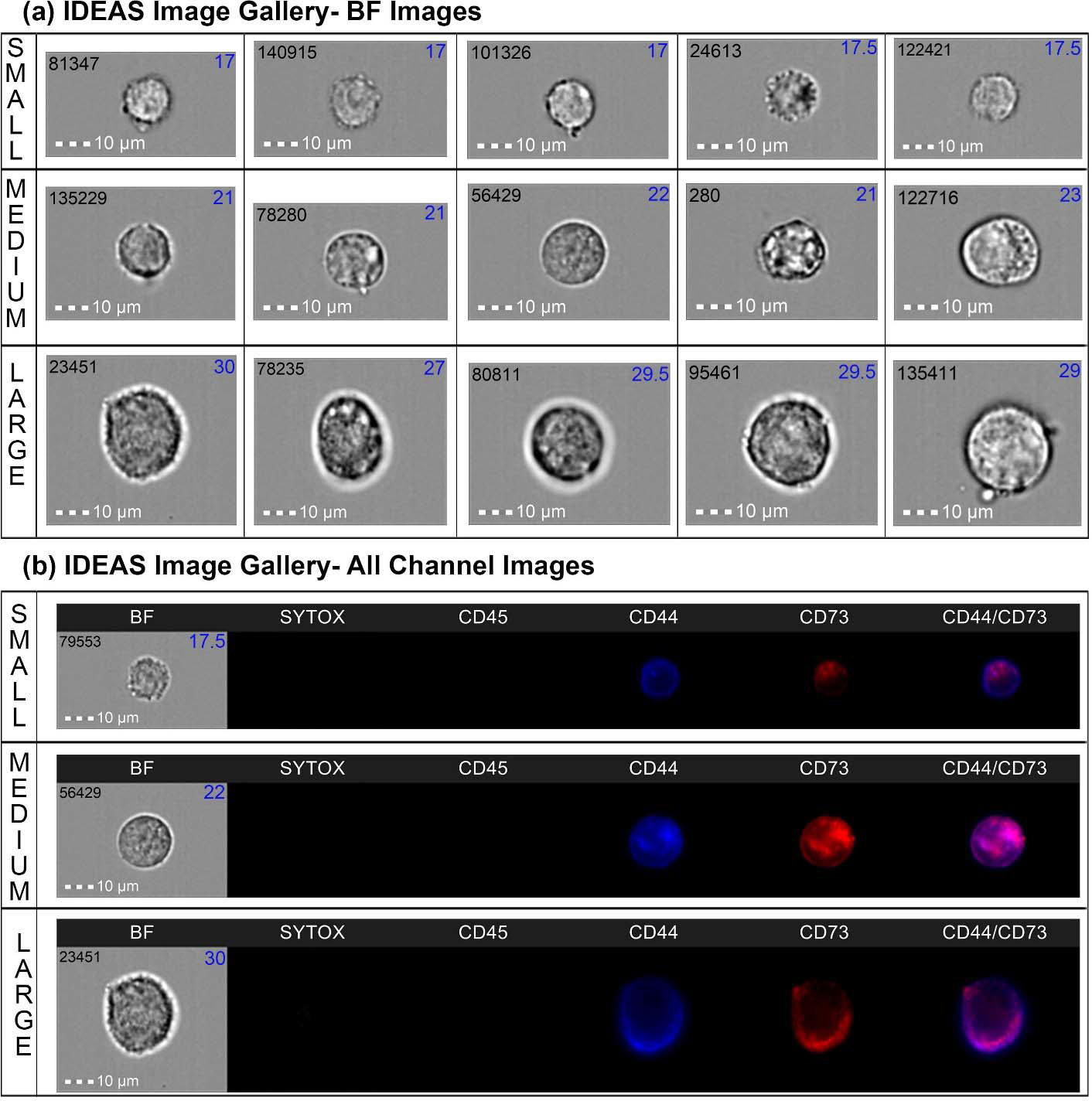
Single cell images shown from the IDEAS image gallery. Each number indicated on the upper left corner is the cell/event number, and the number on the upper right corner (in blue) indicates the width of the cells. **(a)** Brightfield images obtained from IDEAS image gallery show the heterogeneity in sizes, categorized into small, medium, and large. **(b)** Three different categories of cells in all 5 channels along with the colocalization mask of CD44 and CD73 have been shown here.

### Analysis using a novel pipeline-MSCProfiler

We have outlined all the steps that went into building our novel pipeline-the MSCProfiler. The prime benefit of this lay in automation of the analysis workflow, starting from image quality control right up to the parameter estimation and classification of cells, described in Fig. 3(b). Post segmentation, the three features which are extracted from the imaging modalities were the geometric and texture features, and intensity values. The ‘Focused Cells’ (54,356) annotated in our pipeline, generated images of hSHEDs which revealed statistically significant heterogeneous cell populations within a single passage of MSCs (P9). The spread or distribution of area (based on number of pixels in a shape) in the cells, as seen in Fig. 5(a,b), demonstrated 3 different categories of populations which were small (2200-2999 pixels), medium (3001-3999 pixels) and large (4008-4493 pixels) where majority of the cells belonged to the small category. The implication this has on the morphometric parameters was revealed further in the extraction of geometrical features. The shape quantification of the hSHEDs enhanced our understanding of the characterization parameters. With the goal of using the minimum necessary measurement features to characterize a single MSC adequately so that it can be unambiguously classified, we chose the aspect ratio, eccentricity, and compactness of a single cell as our discriminators. The performance of the pipeline in determining the shape measurements depended a lot on how the image objects were pre-processed. The distribution of aspect ratio (ratio of image object height vs width) did not show statistical significance between the 3 categories of cells (small, medium, and large), as seen from Fig. 6(a). Analysis of the annotated pixel values using Kruskal-Wallis multiple comparisons test demonstrated that the small cells were more circular (close to 1.0) than the medium and large cells from the same passage (p-value > 0.99). However, the distribution is negatively skewed in the small cells because the whisker and half-box are longer on the lower side of the median than on the upper side.

**Fig 5:**
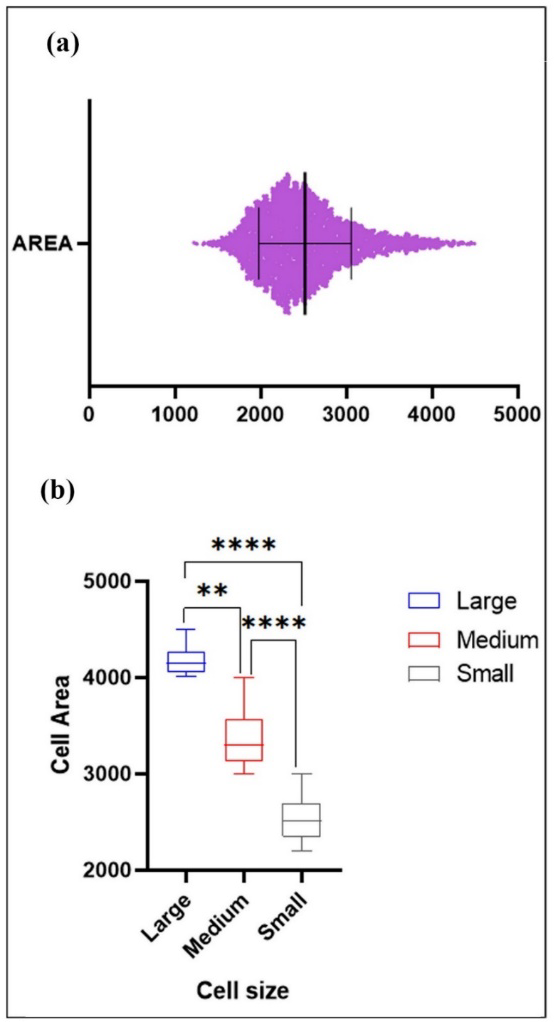
Cell area graphs to demonstrate cell size heterogeneity in hSHEDs using the MSCProfiler. (a) Spread/ distribution of area parameter from cells run through the MSCProfiler pipeline has been graphically represented here. (b) Classification of distinct groups of small, medium, and large cells based on cell area has been shown graphically. Statistical significance was determined using Kruskal-Wallis multiple comparisons test in GraphPad Prism. 3486 cells as an output of the MSCProfiler pipeline were graphically plotted.

**Fig 6:**
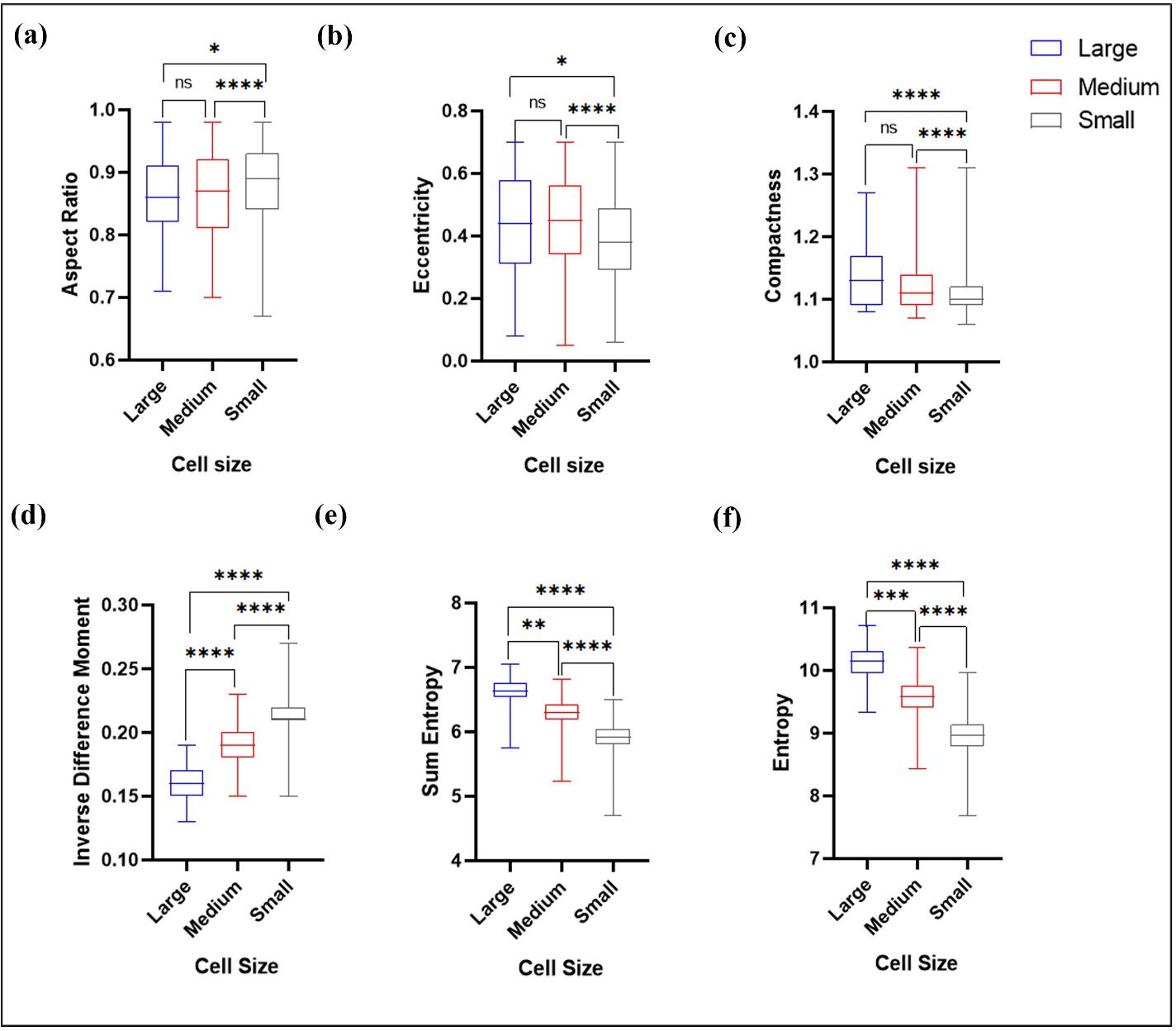
Geometrical and texture parameters demonstrate heterogeneity in hSHEDs using the MSCProfiler. (a-c) Geometrical features such as aspect ratio, eccentricity, and compactness have been compared among the three different classes of cells. (d-f) Texture features such as inverse difference moment, sum entropy and entropy values have been compared between the three different classes of cells. Statistical significance was determined using Kruskal-Wallis multiple comparisons test in GraphPad Prism.

Eccentricity feature (ratio of minor axis length to the major axis length of an image object) distributed the pixel values (between 0-1) and demonstrated that the small cells had lower eccentricity values than the medium and large cells in the same passage, shown in Fig 6(b). The centre of distribution of the box and whisker plots is the lowest of the three distributions in the small cells (median is 0.2-0.4), while in the others the median in between 0.4-0.6. The distribution in all three classes was approximately symmetric, as both the half boxes were almost the same length on both upper and lower sides. According to the statistics, small cells were more circular than the other two categories.

Cell shape measure can be best calculated from descriptor such as mean compactness (ratio of the area of an image object to the area of a circle within the same perimeter). As circle is depicted by a minimum value of 1.0. Larger the compactness more are the irregularities and complexities of the cell boundary. The small cells showed the least compactness values than the medium and large cells. The centre of distribution of the box and whisker plots is the lowest of the three distributions in the small cells (median is 1.0-1.1), while the medium have distributed values between 1.07-1.31 and large cells are 1.08 – 1.27. However, the distribution is positively skewed in the small cells because the whisker and half-box are longer on the upper side of the median than on the lower side.

Texture features are also very important computational feature extraction descriptors. They bring about the values between shapes and individual pixel values. An important measure of variation brought about from the Haralick texture features is inverse difference moment. It is a measure of homogeneity in cells, which gets maximized when neighboring image pixels share same values, i.e., while measuring texture analysis two pixels are considered at a single time, the reference and neighbor pixel. The spatial relationship between the reference and neighbor pixel is calculated to understand the gray level differences. There is a stark difference among the 3 classes of cells in terms of inverse difference moment. The small cells are uniformly homogeneous in texture than the other two classes. It provides high discrimination accuracy for images acquired in motion. This discriminator for local homogeneity is lower in medium and much lesser in large cell types. The centre of distribution of the box and whisker plots is the highest of the three distributions in the small cells (median is 0.2-0.25), while the medium have distributed values close to 0.2 and large cells are close to 0.15.

Entropy of population analysis can reveal highly structured cellular patterns. Image flow cytometry data set distributions have been utilized for identification of malignancy by analyzing the differences in multidimensional distributions of related entropies. Application of these textural entropy investigations to evaluate the homogeneity and randomness of gray values within the cell image are relatively easy to perform and could rely on digital image analysis of label-free or marked live or fixated cells The entropy and combined entropy of the small cells (9.0) were lowest compared to the medium (9.5) and large cell types (10.71). The centre of distribution of the box and whisker plots in all three sets showed homogeneous entropy patterns. More entropy-based patterns could be identified from the medium and large cells, described in Fig 6 (e,f).

In terms of biomarker expression, CD73-PerCP Cy5.5 and CD44-V450 expression levels were also compared in the hSHEDs. CD73 intensity of expression was higher in the large cells compared to the medium and small cells, as seen from Fig. 7(a). We observed that the CD44 expression levels were higher than CD73 in all the three cell types, represented in Fig. 7(b). To compare the distribution pattern of both the markers, we normalized the cell area from which CD44 and CD73 showed expression, by dividing that area by total area of the cell to obtain percentage expression. Results showed that CD44 was expressed over a larger area when compared to CD73, significantly (p-value <0.0001) when analyzed using the Mann Whitney test shown in Fig 7(c).

**Fig 7:**
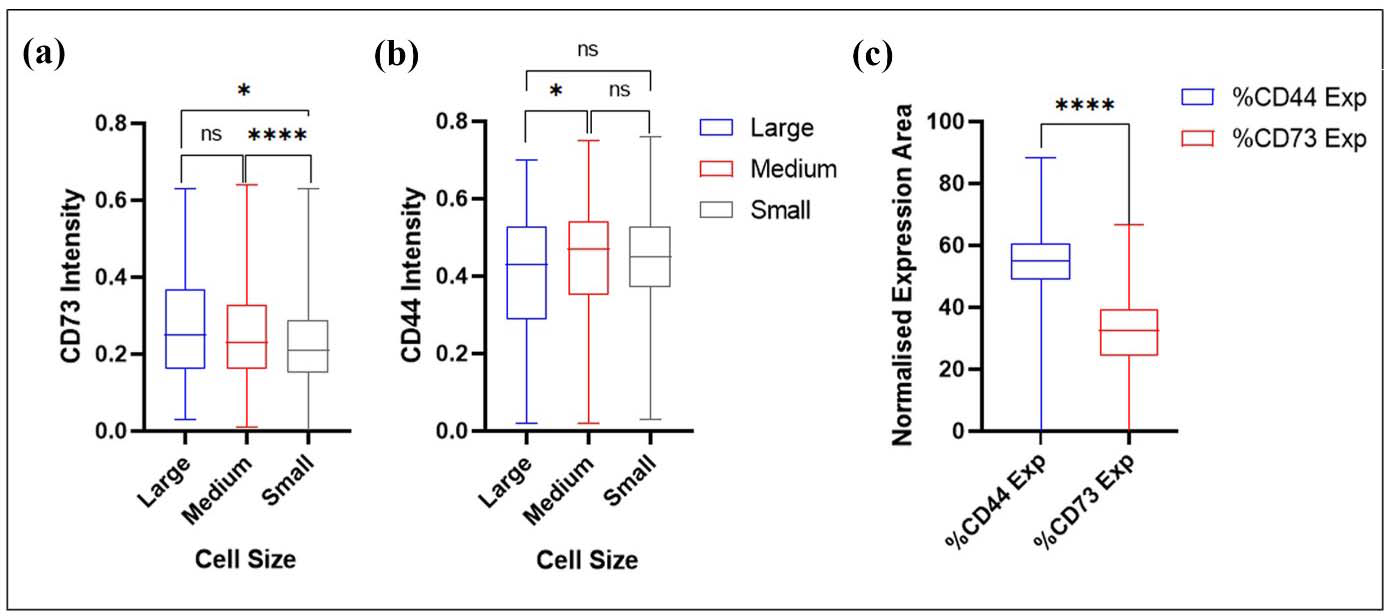
Fluorescence parameters demonstrate heterogeneity in hSHEDs using the MSCProfiler. (a,b) Intensity of CD73 and CD44 expression compared among the three classes of cells. (c) Normalized values of cell area from which markers-CD44 and CD73 express have been compared. CD44 is expressed over a larger area when compared to CD73, significantly (p-value <0.0001). Statistical significance was determined using Mann Whitney test in GraphPad Prism. (‘%CD44 Exp’ and ‘%CD73 Exp’ denote the percentage of CD44 and CD73 expression area, respectively.)

## DISCUSSION

MSC heterogeneity has been documented by many researchers. It can be derived from different sources of tissues or even arise randomly from a clonally dividing cell population. However, what is debatable is whether the appearance of such cellular heterogeneity within the MSC population follows a stochastic or deterministic process. The existence of heterogeneity is proof enough to show that we must not limit our studies at a population level but also look at single cell expression. This will help us come up with more stringent protocols for identifying MSCs best suited for clinical purposes. This brings in the rational of our study in developing an unsupervised pipeline which can assist a stem cell biologist in appropriate identification and classification of MSCs best fit for specific translational studies.

Feature extraction from images of single cells has been the hallmark of this study which led us to develop the MSCProfiler pipeline. The aim was to identify post-acquisition pattern recognition of MSCs (hSHEDs) based on multiple image features of single cells such as aspect ratio, cell texture, shape, surface antigen distribution and intensities of expression of such antigens. We characterized the gross surfaceome of the hSHEDs in this study by developing a non-supervised image processing pipeline that can robustly segment and analyse single-cell morphologies from bright field standalone images. There are multiple robust, high-end screening image analysis software available which can quantify visual cellular morphotypes by microscopy (16). However, the critical steps in any such image analysis tool should be-lack of bias, ability to identify image-based aberrations (image blur, debris crowding, autofluorescence, saturation of pixels) and high speed of resolution of individual image objects. Our pipeline has used images of single cells (hSHEDs) generated from an imaging flow cytometer (Amnis Imagestream Mk II platform) which collectively gives an advantage takes care of most of the image-based aberrations and makes a single image object for acquired data file. This feature rules out the first concern of shadowing of cellular features in case of a smear or a tissue slice (24). We have described an automated protocol implemented in a validated open-source software called CellProfiler (24–26) with a capacity to offer a suite of image-based measurement features which can extract quantitative information from images. Additionally, this software can be useful in leading us to the machine-learning functionality of CellProfiler Analyst for a better user interface.

Although conventional flow cytometry on its own can be a source of high content quantitative tool for all cytometry-related information, IFC has been able to plug in the image data output to further enhance its throughput. We have strategically validated our pipeline with both conventional and image flow cytometry data and demonstrated the shortcomings in either case. Conventional flow cytometry lacks spatial information of every dot on the analysis plot while IDEAS on the Amnis platform is proprietary and needs to be customized to meet individual needs. This has been one of the first attempts to screen for cellular heterogeneity of mesenchymal stem cells using morphometric features of single cells.

## ABBREVIATIONS

AI: Artificial Intelligence BF: Brightfield
.daf: data analysis file
.cif: compensated image file
.ctm: compensation matrix file DF: Darkfield
IFC: Imaging Flow Cytometry
ISCT: International Society for Cell & Gene Therapy
ML: Machine Learning
MSC: Mesenchymal Stem Cell
.rif: raw image file
hSHEDs: human mesenchymal stromal cells obtained from exfoliated deciduous teeth

## Supporting information

Hospital ethical clearance certificate

Institutional stem cell committee approval

Image renaming macro

## ACKNOWLEDGEMENT

We acknowledge the intramural support from Manipal Institute of Regenerative Medicine (MIRM), MAHE Bengaluru Campus, India.

## REFERENCES

1. Andrzejewska A, Lukomska B, Janowski M. Concise Review: Mesenchymal Stem Cells: From Roots to Boost. Stem Cells. 2019 Jul 1;37(7):855–64.

2. da Silva Meirelles L, Chagastelles PC, Nardi NB. Mesenchymal stem cells reside in virtually all post-natal organs and tissues. J Cell Sci. 2006 Jun 1;119(Pt 11):2204–13.

3. Glenn JD. Mesenchymal stem cells: Emerging mechanisms of immunomodulation and therapy. World J Stem Cells. 2014;6(5):526.

4. Saeedi P, Halabian R, Fooladi AAI. A revealing review of mesenchymal stem cells therapy, clinical perspectives and Modification strategies. Stem Cell Investig. 2019 Dec 1;6.

5. Wilson A, Webster A, Genever P. Nomenclature and heterogeneity: Consequences for the use of mesenchymal stem cells in regenerative medicine. Regenerative Med. 2019;14(6):595–611.

6. Dominici M, Le Blanc K, Mueller I, Slaper-Cortenbach I, Marini FC, Krause DS, et al. Minimal criteria for defining multipotent mesenchymal stromal cells. The International Society for Cellular Therapy position statement. Cytotherapy. 2006;8(4):315–7.

7. Pittenger MF, Discher DE, Péault BM, Phinney DG, Hare JM, Caplan AI. Mesenchymal stem cell perspective: cell biology to clinical progress. NPJ Regen Med. 2019 Dec 1;4(1).

8. Laura Elizabeth Lansdowne (2019). Single Cell Analysis – Advantages, Challenges, and Applications | Technology Networks. https://www.technologynetworks.com/drug-discovery/blog/single-cell-analysis-advantages-challenges-and-applications-322768

9. Hennig H, Rees P, Blasi T, Kamentsky L, Hung J, Dao D, et al. An open-source solution for advanced imaging flow cytometry data analysis using machine learning. Methods. 2017 Jan 1;112:201–10.

10. Maguire O, O’Loughlin K, Minderman H. Simultaneous assessment of NF-_κ_B/p65 phosphorylation and nuclear localization using imaging flow cytometry. J Immunol Methods. 2015 Aug 1;423:3–11.

11. Phadwal K, Alegre-Abarrategui J, Watson AS, Pike L, Anbalagan S, Hammond EM, et al. A novel method for autophagy detection in primary cells: Impaired levels of macroautophagy in immunosenescent T cells. Autophagy. 2012 Apr 4;8(4):677.

12. Pugsley HR. Quantifying autophagy: Measuring LC3 puncta and autolysosome formation in cells using multispectral imaging flow cytometry. Methods [Internet]. 2017 Jan 1;112:147–56.

13. Durdik M, Kosik P, Gursky J, Vokalova L, Markova E, Belyaev I. Imaging flow cytometry as a sensitive tool to detect low-dose-induced DNA damage by analyzing 53BP1 and _γ_H2AX foci in human lymphocytes. Cytometry A. 2015 Dec 1;87(12):1070–8.

14. Lee Y, Wang Q, Shuryak I, Brenner DJ, Turner HC. Development of a high-throughput _γ_-H2AX assay based on imaging flow cytometry. Radiation Oncology. 2019 Aug 22;14(1):1–10.

15. Filby A, Perucha E, Summers H, Rees P, Chana P, Heck S, et al. An imaging flow cytometric method for measuring cell division history and molecular symmetry during mitosis. Cytometry A. 2011 Jul; 79(7):496–506.

16. Negm AS, Hassan OA, Kandil AH. A decision support system for Acute Leukaemia classification based on digital microscopic images. Alexandria Engineering Journal. 2018 Dec 1;57(4):2319–32.

17. Helgadottir S, Midtvedt B, Pineda J, Sabirsh A, B. Adiels C, Romeo S, et al. Extracting quantitative biological information from bright-field cell images using deep learning. Biophys Rev. 2021 Sep;2(3):031401.

18. Imboden S, Liu X, Lee BS, Payne MC, Hsieh CJ, Lin NYC. Investigating heterogeneities of live mesenchymal stromal cells using AI-based label-free imaging. Sci Rep. 2021 Dec 1;11(1).

19. Otesteanu CF, Ugrinic M, Holzner G, Chang YT, Fassnacht C, Guenova E, et al. A weakly supervised deep learning approach for label-free imaging flow-cytometry-based blood diagnostics. Cell Reports Methods. 2021 Oct 25;1(6).

20. Sanz G, Manuel Martínez-Aranda L, Tesch PA, Fernandez-Gonzalo R, Tommy X, Lundberg R. INNOVATIVE METHODOLOGY Muscle2View, a CellProfiler pipeline for detection of the capillary-to-muscle fiber interface and high-content quantification of fiber type-specific histology. J Appl Physiol. 2019; 127:1698–709.

21. Zhu H, Mitsuhashi N, Klein A, Barsky LW, Weinberg K, Barr ML, et al. The role of the hyaluronan receptor CD44 in mesenchymal stem cell migration in the extracellular matrix. Stem Cells. 2006 Apr 1;24(4):928–35.

22. Tan K, Zhu H, Zhang J, Ouyang W, Tang J, Zhang Y, et al. CD73 expression on mesenchymal stem cells dictates the reparative properties via its anti-inflammatory activity. Stem Cells Int. 2019;2019.

23. Chan DK, Miskimins WK. Metformin and phenethyl isothiocyanate combined treatment in vitro is cytotoxic to ovarian cancer cultures. J Ovarian Res. 2012;5(1).

24. Bray MA, Carpenter AE. Quality Control for High-Throughput Imaging Experiments Using Machine Learning in Cellprofiler. In: Methods in Molecular Biology. Humana Press Inc.; 2018. p. 89–112.

25. Lamprecht MR, Sabatini DM, Carpenter AE. CellProfiler^TM^: Free, versatile software for automated biological image analysis. Biotechniques. 2007 Jan;42(1):71–5.

26. Bray MA, Vokes MS, Carpenter AE. Using Cellprofiler for automatic identification and measurement of biological objects in images. Curr Protoc Mol Biol. 2015; 2015:14.17.1-14.17.13.

