## Supplementary material for "MSCProfiler: An image processing workflow to investigate Mesenchymal Stem Cell heterogeneity using imaging flow cytometry data": Hospital ethical clearance certificate

SRI
RAJIV GANDHI
COLLEGE OF DENTAL SCIENCES & HOSPITAL

Recognised by Dental Council of India, New Delhi
Affiliated to Rajiv Gandhi University of Health Sciences, Karnataka

RGC Campus, Cholanagar, R.T. Nagar Post, Bangalore – 560 032, Karnataka – INDIA


www.srgcds.ac.in

No. SRGCDS/2020/374

### ETHICAL COMMITTEE

| | |
|---|---|
| Name of investigator | Uttara Chakraborty Ph.D., Assistant Professor, Manipal Institute of Regenerative Medicine (MIRM), Bangalore- 560065 |
| Study Title | To study the crosstalk of dental pulp stromal cells with the commensal fungi of the oral cavity. |
| Date of approval | 30/12/2020 |

The Ethical Committee, Sri Rajiv Gandhi College of Dental Sciences, has reviewed and approved your study.

| S/No | Name & Designation | Position | Signature |
|---|---|---|---|
| 1 | Dr. Silju Mathew,Assoc.Dean,FDS- RUAS | Chairman | [signature] |
| 2 | Dr. Tejavathi Nagaraj, Prof & HoD - Oral Medicine & Radiology | Member | [signature] |
| 3 | Dr. Umesh Yadalam, Prof & HoD - Periodontology | Member | [signature] |
| 4 | Dr. Akshay D. Shetty, Prof & HoD - Oral & Maxillofacial Surgery | Member | [signature] |
| 5 | Dr. Sushant Anant Pai, Principal Prof & HoD - Prosthodontics | Member | [signature] |
| 6 | Dr. Rajesh R.N.G , Prof & HoD - Orthodontics | Member | [signature] |
| 7 | Dr. Kusum Valli. S, Prof & HoD - Conservative Dentistry | Member | [signature] |
| 8 | Dr. Yogesh T.L , Prof & HoD - Oral Pathology | Member | [signature] |
| 9 | Dr. Santhosh T. Paul, Prof & HoD - Pedodontics | Member | Santhosh |
| 10 | Dr. Sida Tagore, Asso. Prof - General Pathology | Member | Sida Tagore |
| 11 | Dr. B.P.P. Rathnam, Professor - Anatomy | Member | B Rathnam |
| 12 | Mr. Sunil. S - Legal Advisor | Member | [signature] |
| 13 | Mr. G. A Bawa - NGO Representative | Member | [signature] |
| 14 | Mr. Jayappa Reddy - Social Worker | Member | [signature] |

The Ethical Committee, SRGCDS expect to be informed about any changes in the protocol and patient information concerned and asked to be provided a copy of the final report.
