## Supplementary material for "MSCProfiler: An image processing workflow to investigate Mesenchymal Stem Cell heterogeneity using imaging flow cytometry data": Institutional stem cell committee approval

### MANIPAL INSTITUTE OF REGENERATIVE MEDICINE

BENGALURU

*(A constituent unit of MAHE, Manipal)*

INSPIRED BY LIFE

#### Institutional Committee for Stem Cell Research (IC-SCR), Manipal Institute of Regenerative Medicine (MIRM), Manipal Academy of Higher Education (MAHE), Bangalore-560 065

Date: 16/08/2021

To
Dr. Uttara Chakraborty
Principal Investigator

**Sub: Project proposal No: MIRM-ICSCR/004/2021**

The Committee has reviewed the proposal titled "Investigation of immunophenotypic heterogeneity and lineage specific differentiation potential in oral mesenchymal stromal cells" and approved this project in its meeting held on July 15th, 2021 for a period of 3 years.

The following members of the IC-SCR were present at the meeting held on 15/07/2021 at 2:15 PM, at Manipal Institute of Regenerative Medicine, Manipal Academy of Higher Education (MAHE), Bangalore-65.

| | |
|---|---|
| Prof. M.R.S Rao | Chairperson |
| Prof. Sudhir Krishna | Vice-Chairperson/ Stem Cell Expert |
| Mr. Vasantha Krishna K | Legal Expert |
| Dr. Sreevatsa | Ethics Expert |
| Dr. Abha Rao | Social Scientist |
| Dr. Monojit Debnath | Geneticist / Cell and Molecular Biology Expert |
| Dr. Vijay Bhat | Biochemist |
| Dr. Shrikrishna Isloor | Veterinarian |
| Dr. Kruti Varshney | Medical Expert |
| Mrs. C P Yashoda | Layman |
| Dr. Manasa Nune | Member Secretary |

[illegible]

Signature of the Chairman

Copy forwarded to protocol applicant
Copy in file
