## Supplementary material for "MSCProfiler: An image processing workflow to investigate Mesenchymal Stem Cell heterogeneity using imaging flow cytometry data": Image renaming macro

//*********************************************************************************************************************************************************************
/// 	Program: This script is to rename and save as a new tif file
///		Author: Dr. Lakshmi Balasubramanian
///		Usage: Using the images that are extracted using ideas with .ome.tif this script save as .tif files that is readable in all tools
///		Input: Folder that contains images with ".ome.tif" files; Also choose the output folder where you want to save the new renamed images
///		Output: User can choose the output folder, in which the results are stored. 
///		Other: The line with the comments starts with "//" in which description and intructions are given.
///		So if a line starts witgh "//" the program will not execute that line.
//*******************************************************************************************************************************************************************

//Turn on the Batch mode


setBatchMode(true);

//Getting the input and output folder from the user
dir=getDirectory("Choose an Input Directory"); 
//print(dir); 
list = getFileList(dir); 
saveloc = getDirectory("Choose Ouput Directory");
//print(saveloc);


//Looping through all the files in the input folder
for (i=0; i<list.length; i++){
	open(dir+list[i]);
	
	filename = dir+list[i];
    //print(filename);
    
    
	Orig_imgName=getTitle();
	print(Orig_imgName);
	selectWindow(Orig_imgName);

	New_imgName=replace( Orig_imgName , ".ome.tif" , ".tif" );
    //print(New_imgName);
	
	saveAs("Tiff", saveloc + New_imgName);

	//Closing all the windows opened
	run("Close All");
}

waitForUser("Done successfully!!");
